# EvolvingSTEM: A microbial evolution-in-action curriculum that enhances learning of evolutionary biology and biotechnology

**DOI:** 10.1101/514513

**Authors:** Vaughn S. Cooper, Taylor M. Warren, Abigail M. Matela, Michael Handwork, Shani Scarponi

## Abstract

Evolution is a central, unifying theory for all of life science, yet the subject is poorly represented in most secondary-school biology courses, especially in the United States. One challenge to learning evolution is that it is taught as a conceptual, retrospective subject with few tangible outcomes for students. These typical passive learning strategies lead to student disengagement with the material and misunderstanding of evolutionary concepts. To promote greater investment and comprehension, we developed EvolvingSTEM, an inquiry-based laboratory curriculum that demonstrates concepts of natural selection, heredity, and ecological diversity through experimental evolution of a benign bacterium. Students transfer populations of *Pseudomonas fluorescens* growing on plastic beads, which selects for biofilm formation and mutants with new, conspicuous phenotypes. We introduced our curriculum to four introductory high school biology classes alongside their standard curriculum materials and found that students who learned evolution through EvolvingSTEM scored significantly better on a common assessment targeted to Next Generation Science Standards than students taught only the standard curriculum. This latter group subsequently achieved similar scores once they too completed our curriculum. Our work demonstrates that inquiry-based, hands-on experiences with evolving bacterial populations can greatly enhance student learning of evolutionary concepts.

## Introduction

Understanding evolutionary processes is fundamental to all areas of life science because evolution serves as a conceptual framework to organize other life science topics, such as organismal diversity and ecological interactions. Furthermore, some of the most significant threats to human health are evolutionary phenomena; therefore, knowledge of evolutionary processes has a direct impact on public health and medicine (Wells et al. 2017). For example, antimicrobial resistance and cancer are caused by the rapid evolution of microbes and our own cells, respectively (Karatan and Watnick 2009; Greaves and Maley 2012; Berendonk et al. 2015; Makohon-Moore and Iacobuzio-Donahue 2016; Alizon and Méthot 2018). In addition, ongoing revolutions in biotechnology and personalized medicine, such as gene-editing (i.e., CRISPR), can only be understood in the context of the evolutionary concept of descent from a shared ancestral lineage (Makarova et al. 2015; Knott and Doudna 2018). A strong knowledge base of evolution is therefore invaluable for a literate society to understand scientific and medical advances and for a prepared workforce to excel in jobs in science, technology, and engineering. The value of evolutionary biology knowledge is highlighted by its inclusion as a core concept for STEM education practices (National Research Council 2012; NGSS Lead States 2013; NSTA 2013).

Although the importance of evolutionary biology is well-established, misconceptions of its basic principles remain prevalent among students, the general public, and even the teachers who are providing instruction (Cunningham and Wescott 2009; Gregory 2009; Sickel and Friedrichsen 2013; Yates and Marek 2014; Glaze and Goldston 2015). While many concurrent factors likely contribute to poor understanding (Smith 2010a; 2010b; Pobiner 2016), one potential reason that evolutionary concepts are misunderstood is that typical curricula use passive learning strategies, where instruction relies on lectures and textbook readings. Current evolution curriculum design runs counter to evidence that student-centered, active learning strategies are the most effective method for science teaching and have been shown to improve student understanding of evolutionary concepts (Nehm and Reilly 2007; Nelson 2008; Freeman et al. 2014; Romine et al. 2017). Courses that provide students with authentic research experiences are especially effective at increasing student engagement and promoting a deeper understanding of evolution (Jordan et al. 2014; Ratcliff et al. 2014; Broder et al. 2018).

There is therefore a critical need for engaging and informative evolutionary biology curricula that provide K-12 students the opportunity to explore the concept of changing frequencies of inherited traits just as they attempt to quantify gravity in physics or acid-base reactions in chemistry. To meet this need, we developed EvolvingSTEM, a curriculum that provides inquiry-based learning of evolution, microbiology, ecology, and heredity with a laboratory experiment that employs real scientific research practices. EvolvingSTEM allows students to visualize evolutionary adaptations arising in real time by growing populations of the harmless bacterium *Pseudomonas fluorescens* under conditions that select for the formation of a biofilm. A biofilm is a surface dwelling community of microbes encased in a protective coating of self-produced polymers; biofilms are the dominant form of microbial life (Costerton et al. 1987). They are also structured, heterogeneous environments that include varied ecological niches (Karatan and Watnick 2009). Bacteria with advantageous mutations colonize these niches, and their adaptations cause visible differences in colony morphology from the ancestral genotype (Rainey and Travisano 1998; Flynn et al. 2016). This evolution-in-action occurs within days, requires little specialized equipment, and can be offered in any classroom laboratory that can support sterile technique. Our curriculum is intended to replace standard, passive learning curricula to meet competencies for natural selection and evolution described in the Next-Generation Science Standards (HS-LS4, (NGSS Lead States 2013)). We hypothesized that students who learn evolutionary concepts with our curriculum would have significant increases in content knowledge relative to students that were provided only the standard curriculum.

## Results

### Developing and refining an amenable protocol for teaching bacterial evolution to high school students

The idea to teach evolutionary concepts to high school students with a bacterial evolution experiment grew from our research on identifying the causes of rapidly evolving mutant colony morphologies of the opportunistic pathogens *Burkholderia cenocepacia* and *Pseudomonas aeruginosa* (Poltak and Cooper 2011; Flynn et al. 2016). These species are particularly threatening to persons with cystic fibrosis, where they cause chronic airway infections by forming biofilms (Starkey et al. 2009; Ashish et al. 2013). Biofilm-associated infections are inherently more resistant to host immunity and antimicrobials because secreted adhesive polymers are protective and the cells within grow more slowly (Harrison et al. 2005). Eventually, some bacteria disperse from the colony, either as individuals or clusters, to inhabit new surfaces and resume the biofilm lifecycle (Poltak and Cooper 2011; Martin et al. 2016).

In order to study the dynamics of bacterial evolution *in vitro*, we developed a simple method to model the biofilm lifecycle of surface attachment, biofilm formation, dispersal, and recolonization (Figure 1, (Poltak and Cooper 2011; Traverse et al. 2013; O’Rourke et al. 2015; Flynn et al. 2016; Turner et al. 2018). In short, we culture bacteria for 24 hours in test tubes containing growth media and a polystyrene bead. A subset of the bacteria colonize the bead and form a biofilm. We then transfer only the biofilm-covered bead to a new tube with a fresh bead. We repeat this process daily to select for bacterial mutants that are best adapted to aspects of the entire biofilm lifecycle. Conveniently, we found that biofilm adapted mutants also display altered colony morphologies when grown on agar plates, making them conspicuous to students.

**Fig. 1.**
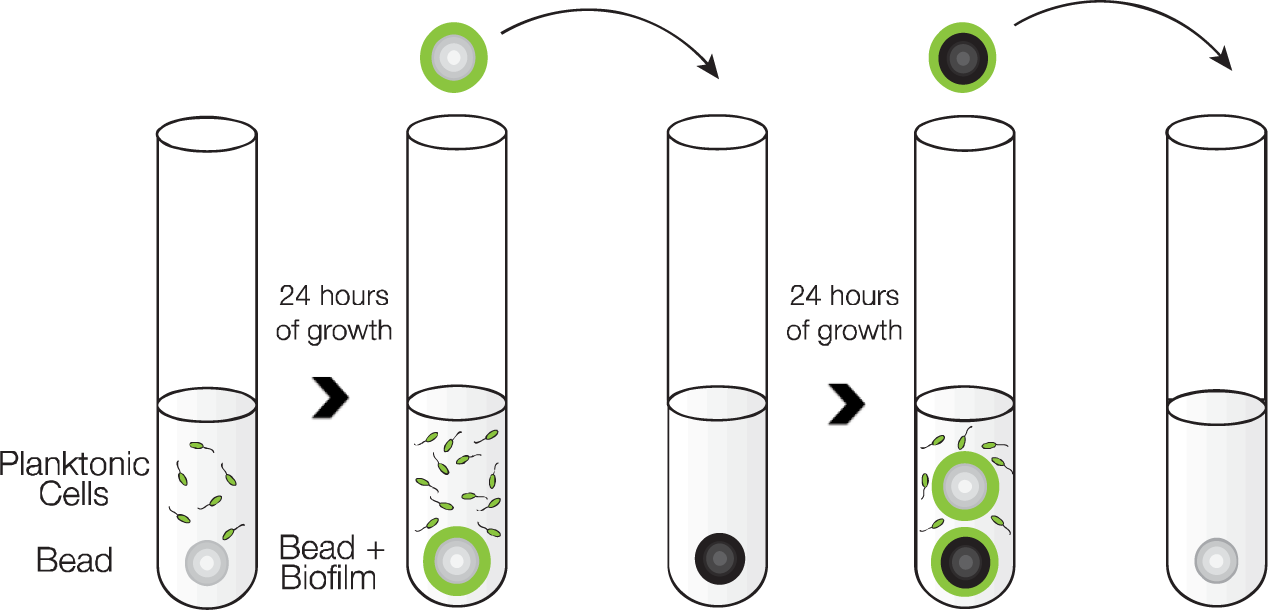
Biofilm lifecycle model. Bacteria are grown in test tubes with plastic beads on which biofilm forms. Daily bead transfers select for bacterial attachment, assembly, dispersal, and reattachment. Figure adapted from (Turner et al. 2018).

In collaboration with science teachers and administrators at Winnacunnet High School (Hampton, NH), we modified our research laboratory protocol to accommodate implementation in a high school classroom. We selected the plant probiotic bacterium, *Pseudomonas fluorescens* SBW25, as our study subject because it had several qualities that made it a good candidate for use in a high school classroom: (1) it is benign, and thus safe for students with no microbiology experience, (2) it had previously been suggested as a good candidate for use in educational settings (Green et al. 2011; Spiers 2014), and (3) it is the subject of a large body of research on its capacity for rapid and conspicuous adaptive evolution in biofilm-related conditions (Rainey and Travisano 1998; Spiers 2005). Adaptive *P. fluorescens* mutants are often characterized by rugose or rosette-like colony morphologies resulting from greater production of polysaccharides for attachment (Rainey et al. 2000). We found that experimental evolution of *P. fluorescens* SBW25 in the biofilm lifecycle model selected for a high frequency of adaptive mutants with novel colony morphologies in less than two weeks.

To accelerate this process and ensure that our experiment could be performed within the timeframe of a high school biology lesson, we conducted a series of trials in different media to determine conditions that resulted in predictable, rapid adaptations. We found that growth in King’s B medium (KB) generated multiple, heritable colony phenotypes within seven days. In the interest of accelerating the evolutionary dynamics, we repeated the experiment in KB medium with various glycerol concentrations. We found that an increase from 1.5% to 2.5% glycerol selected for novel colony morphologies at detectable frequencies in four days. We named this modified media recipe “Queen’s B” (QB) and used this recipe thereafter. Media recipes are available in a supplemental file (Supplemental File 1).

Students can use our modified protocol to guide an inquiry-based experiment that allows them to visualize evolution in their bacterial populations in only six class periods (Fig 2). For example, on Monday, students inoculate glass test tubes containing QB media and a polystyrene bead with a clone of *P. fluorescens* SBW25, and then perform bead transfers for the following three days (Tuesday-Thursday). During the process of bead transfer, students can identify effects of natural selection by observing increased biofilm production on the walls of their test tubes. In addition, at the beginning and end of the week, students sample their populations by growing individual bacterial colonies on agar plates. Students can make observations of mutant colonies on the Monday of the following week and compare these colonies to those of the ancestral population that were plated earlier in the week. Students can be given additional curriculum materials, such as homework and pretests, to prepare them for each step in the laboratory protocol and provide opportunities for them to link the heritable, adaptive evolutionary change they observe in their experiment to the evolutionary processes that produced this dynamic. Through EvolvingSTEM, students can acquire the knowledge to meet Next Generation Science Standards for Natural Selection and Evolution (Box 1; (NGSS Lead States 2013)). Curriculum materials are available as supplemental files (Supplemental Files 2-4).

**Fig. 2.**
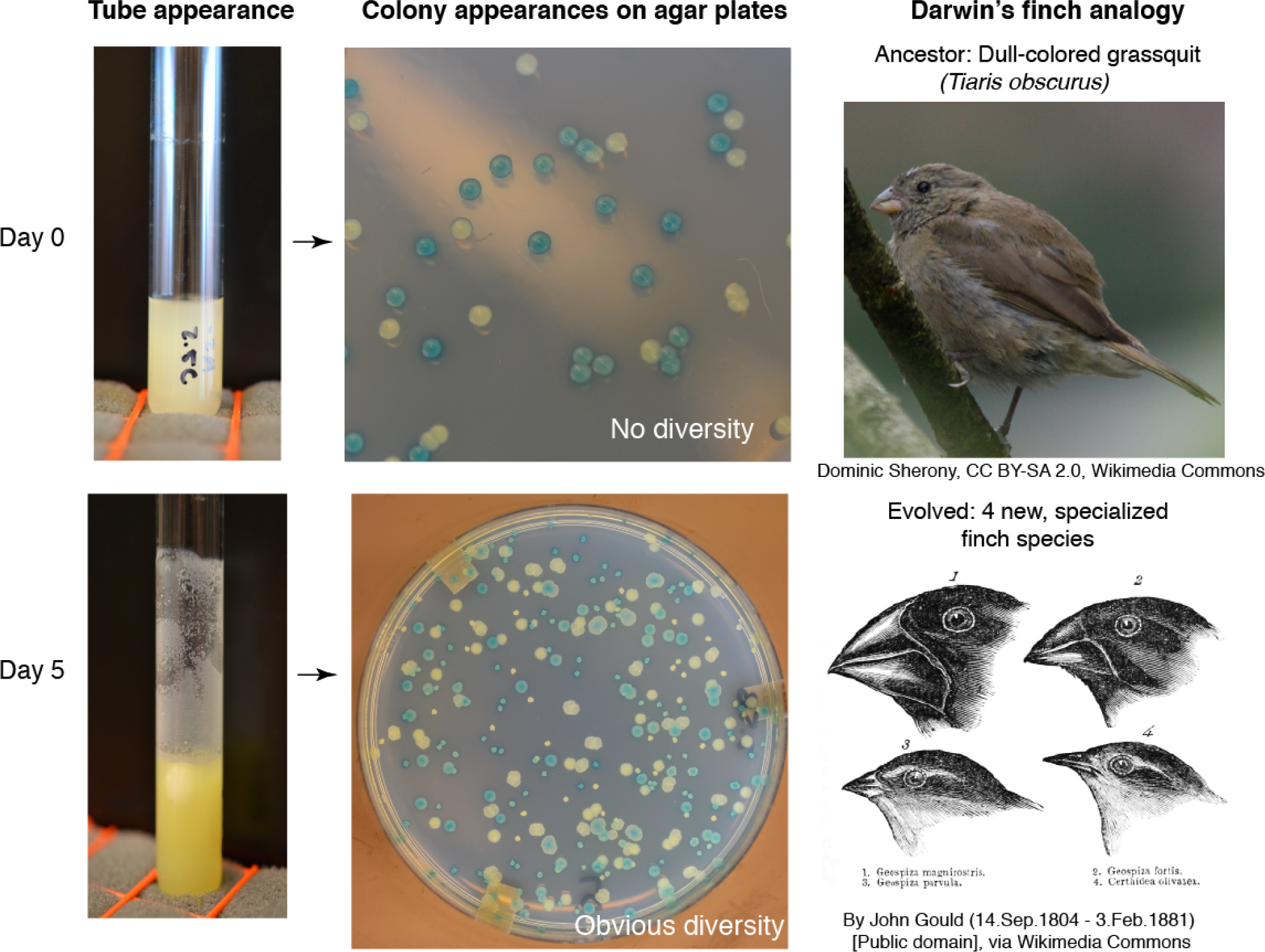
Adaptation to biofilm selection can occur within days and produce conspicuous phenotypic differences. Populations were founded with equal ratios of Lac+ (blue) and Lac-(white) ancestral genotypes that do not differ in morphology. After 5-7 days, new colony morphologies evolve and represent different biofilm-associated ecological strategies, as different beak shapes of Darwin’s finches represent distinct feeding strategies (Rainey and Travisano 1998, Poltak and Cooper 2011).

### Learning outcomes

The exact outcome of any individual experiment is unknown because the biofilm selection acts on randomly occurring mutations in the bacterial populations that were founded from a single clone. In fact, this variability among these independent “replays” of evolution is realistic and demonstrates effects of chance and contingency on evolution (Blount et al. 2018). Nonetheless, student groups propagate multiple populations in different culture tubes under identical experimental conditions, and this replication means they are very likely to see mutants with novel morphologies in at least one experimental population. In addition, students compare their experimental populations to a control population that does not contain the bead and therefore is not under selection for increased biofilm production. Students can examine the phenotypes found in each population over time, compare their findings to those of other classmates, and develop their own explanations for their observations. This allows students to apply the comparative method of evolutionary biology and begin the process of scientific inquiry. Students are encouraged to consider why their replicate populations vary and propose reasons for that variation, ranging from experimental error, to peculiarities of the bead transfers, to genuine evolutionary randomness.

**Box 1.** Next Generation Science Standards (NGSS) Targeted by EvolvingSTEM. NGSS (2013) are based on *A Framework for K-12 Science Education: Practices, Crosscutting Concepts, and Core Ideas* (National Research Council 2012) and designed through a collaboration between 26 states, the National Research Council, the National Science Teachers Association, the American Association for the Advancement of Science, and Achieve, Inc.

EvolvingSTEM provides students with the knowledge to meet the following NGSS HS-LS4 standards. These are performance expectations.

1. Communicate scientific information that common ancestry and biological evolution are supported by multiple lines of empirical evidence.
2. Construct an explanation based on evidence that the process of evolution primarily results from four factors: (1) the potential for a species to increase in number, (2) the heritable genetic variation of individuals in a species due to mutation and sexual reproduction, (3) competition for limited resources, and (4) the proliferation of those organisms that are better able to survive and reproduce in the environment.
3. Apply concepts of statistics and probability to support explanations that organisms with an advantageous heritable trait tend to increase in proportion to organisms lacking this trait.
4. Construct an explanation based on evidence for how natural selection leads to adaptation of populations.
5. Evaluate the evidence supporting claims that changes in environmental conditions may result in (1) increases in the number of individuals of some species, (2) the emergence of new species over time, and (3) the extinction of other species.

In addition, for HS-LS4-2, students will learn:

- Random mutation results in genetic variation between members of a population.
- Genetic variation can result in trait variation that leads to performance differences among individuals.
- Competition for limited resources results in differential survival. Individuals with more favorable phenotypes are more likely to survive and reproduce, thus passing traits to subsequent generations.
- Evolutionary fitness is measured by reproductive success.
- An adaptation is a heritable genetic variant manifested as a trait that provides an advantage to an individual in a particular environment.
- In addition to natural selection, chance and random events can influence the evolutionary process, especially for small populations.

In addition, students will be skilled at:

- Developing experimental investigations that can be used to test specific hypotheses.
- Evaluating evidence to qualitatively and quantitatively investigate the role of natural selection in evolution.
- Constructing evidence-based explanations that the process of evolution is a consequence of the interaction of four factors: (1) the potential for population size to increase, (2) genetic variation, (3) competition for resources, and (4) proliferation of individuals better able to survive and reproduce in a particular environment.
- Applying basic mathematics to calculate the fitness advantages of selected mutants and/or to compare differences in levels of biofilm production.
- Developing generalizations of the results obtained and/or the experimental design and applying them to new problems, including the design of new experiments and interpreting results in the context of natural and infectious bacterial biofilms.

The speed of adaptation in biofilm models results from strong selection for more adherent mutants that bind not only the provided surface (e.g. polystyrene), but also other attached bacteria or secreted substances. Consequently, selection often favors the evolution of diverse, conspicuous phenotypes within each tube and not just a single, more adherent type. This result not only simulates the process of adaptive radiation often illustrated using Darwin’s finches in textbooks (Figure 2), but also reproduces the selection for traits associated with adherence that often occurs during biofilm-associated infections (Traverse et al. 2013; Cooper et al. 2014; O’Rourke et al. 2015; Gloag et al. 2018). The “wrinkly” colony morphologies that evolve in our model are genetically and functionally identical to those commonly isolated from infections of the related species *Pseudomonas aeruginosa* in the airways of cystic fibrosis patients and in chronic skin wounds (Starkey et al. 2009; Gloag et al. 2018). Students can therefore connect their classroom experiments to recent findings at the interface of evolutionary biology and medicine to see how basic biological research impacts their everyday lives. Furthermore, making connections from classroom activities to real-world examples can increase students’ understanding of evolution and their engagement with the material (Beardsley et al. 2011; Infanti and Wiles 2014).

### Assessment of student learning

We used a delayed intervention approach to assess learning in 4 classes of 9^th^ grade biology honors students at Winnacunnet High School, a suburban public high school in New England. Group 1 included classroom A, taught by MH, and classroom B, taught by SS. This group used an earlier version of our EvolvingSTEM curriculum that did not use a control population alongside their standard curriculum materials, which included textbook readings, lectures, and an educational video. Group 2 included classrooms C and D, both taught by SS. This group first received the standard curriculum with additional lecture materials, followed by EvolvingSTEM (Table 1). Students conducted the experiments and analyses for our curriculum in groups of three or four individuals, requiring collaborative teamwork.

**Table 1:** Composition of Study Groups.

| Group | Class – Teacher | Number of Students per Class | Total Number of Students per Group |
| --- | --- | --- | --- |
| 1 | A – Teacher MH | 19 | 41 |
|  | B – Teacher SS | 22 |  |
| 2 | C – Teacher SS | 18 | 37 |
|  | D – Teacher SS | 19 |  |

A summative assessment was used to determine whether students achieved an increased understanding of evolutionary concepts. The test consisted of multiple choice and free response questions to address student learning of higher-order critical thinking aligned to NGSS. Specifically, test questions were devised to assess whether students met NGSS (2013) performance expectations HS-LS4-1,2,3, and 5. We developed a grading rubric for the free response questions based on templates suggested by Wiggins and McTighe (2005) that required answers with accurate information, specific vocabulary, and a well-structured defense that incorporated outside examples (Wiggins and McTighe 2005). Our assessment and grading rubric are available as supplemental files (Supplemental File 5). All assessments were conducted by one of us (TW) on anonymized tests as proscribed by our IRB.

Pretests were given to both groups prior to the start of classroom evolution activities. Group 1 students were given a posttest after completing the EvolvingSTEM curriculum. Group 2 students were given a midtest after completing the standard curriculum, and then a posttest after completing EvolvingSTEM. We found no significant difference between the average pretest score of Group 1 and Group 2 students (13.17 (26%) vs. 12.5 (25%) out of 50 points total; t=0.60, p=n.s.), indicating that all students began with a similar knowledge base (Fig. 3). Quantitative analyses of student knowledge gains revealed that students who completed EvolvingSTEM (Group 1) showed significant improvement on their average posttest scores, with an average gain of 19.16 points, thereby increasing their overall score by 38% between the pre- and posttest (t=16.61, p<0.0001). Students provided the standard curriculum (Group 2) also showed significant improvement on their average midtest score, which increased by 10.14 points (t= 9.72, p<0.0001), resulting in an overall increase of 21% between pre-and midtest. Although both student groups showed improvement, Group 1 achieved significantly higher average test scores after completing EvolvingSTEM than Group 2 did after completing the standard curriculum (t=5.87, p<0.0001). Students who learned evolution with EvolvingSTEM therefore achieved significantly greater gains in comprehension of evolution than students who learned it from the standard curriculum.

**Fig. 3.**
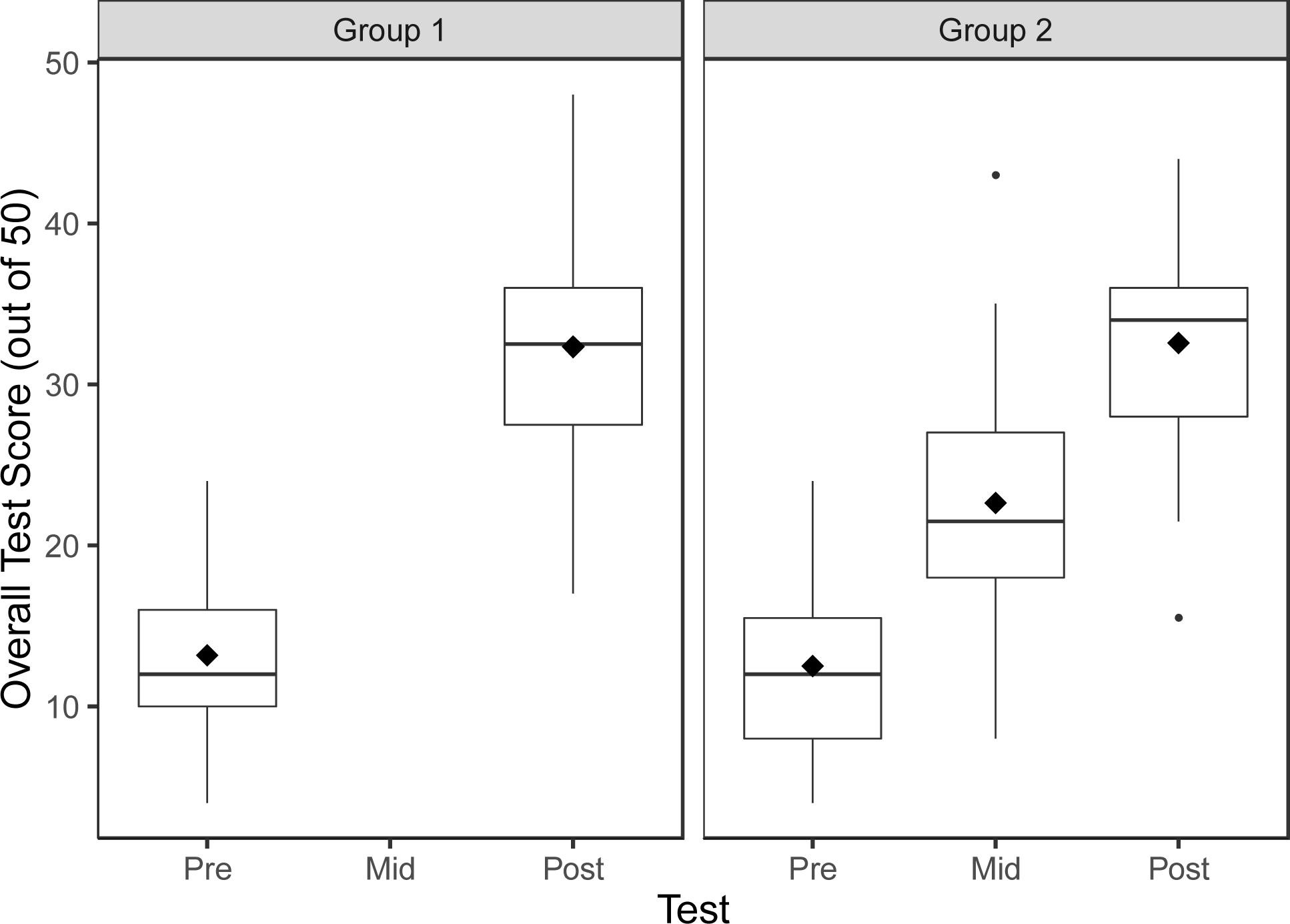
Boxplot of student assessment scores. The EvolvingSTEM curriculum produces significantly greater gains in comprehension of NGSS topic HS-LS-4 than the standard curriculum (Group 1 Post vs Group 2 Mid, t=5.87, p<0.0001). After experiencing our curriculum, Group 2 students subsequently achieved equivalent scores to Group 1 students (Group 1 Post vs Group 2 Post, t=0.14, ns). Mean values are indicated with diamonds.

Once students in Group 2 were exposed to EvolvingSTEM, their average posttest scores had an overall increase of 20% in comparison to their midtest scores, reaching knowledge gains made by Group 1 students (Fig. 3). Knowledge gains by both Groups were overwhelmingly attributable to increased scores on the free-response section of the assessment. Average free-response scores from pretests to posttests increased by 18.09 points (48%) for Group 1 students and 20.59 points (54%) for Group 2 students. In comparison, average multiple-choice scores increased by 1.07 points for Group 1 students and decreased by 0.54 points for Group 2 students. These results may indicate that EvolvingSTEM has a greater impact on improving students’ higher-order cognitive skills, such as applying knowledge to an unknown problem and performing data analysis. There was no significant difference between Group 1 and 2 posttest scores (t=0.14, p=n.s.), even though Group 2 students were provided more detailed verbal instruction and took one additional assessment. This result speaks to the power of EvolvingSTEM to increase student knowledge and suggests that our curriculum can serve to replace, rather than supplement, the standard evolution curriculum.

## Discussion

We developed an inquiry-based microbiology curriculum to improve the engagement of high school biology students with topics central to evolutionary biology and their subsequent understanding of related NGSS concepts. We observed high levels of engagement when students participated in our curriculum. Students were assigned concept and readiness tests each night to ensure that they arrived prepared for the next day’s microbiology experiments and evolution curriculum. Their high rates of completion indicated increased enthusiasm. While we acknowledge this is a simple observation, teachers and coauthors (MH and SS) also indicated that students who rarely participated in class-based discussions emerged as enthusiastic group leaders while performing the EvolvingSTEM experiment. Informal post-surveys of student attitudes towards the curriculum were overwhelmingly positive. Students indicated that they were enthusiastic about the bacterial model, enjoyed coming to class to work on the experiment, and felt that our curriculum was better at teaching them than the standard lecture-style class. The group format for the experiments and analyses encouraged the students to collaborate and support one another throughout the program. Students tended to hold one another accountable, but also demonstrated cohesion when groups compared their replicate populations, demonstrating both friendly competition and pride and ownership in their results. Further, many students expressed that they felt like “real scientists” using equipment like pipettes, vortexes, and the incubator. They shared a greater sense of what science was actually like and asked more questions about microbiology and evolution research and other scientific careers.

Crucially, teachers found EvolvingSTEM to be effective at demonstrating evolution in action, thereby increasing student understanding of natural selection, mutation, and the effects of chance, and increasing student interest and engagement with biology. Student assessments also demonstrated the substantial benefit of our curriculum to student learning, and consequently, our curriculum replaced the standard, honors biology WHS evolution curriculum in subsequent years. The sustainability of the EvolvingSTEM curriculum has been greatly facilitated by the involvement of returning students who demonstrated particular interest in the program and who served as *de facto* teaching assistants through an Extended Learning Opportunity program. (More information about this program will be the subject of a future report.) This teaching experience was made possible by engaging first-year students in laboratory research, which allowed them to help teach new students for up to three subsequent years prior to graduating.

We found that EvolvingSTEM provided students with significant learning benefits in comparison to standard curricula. After completing our curriculum, students achieved significantly higher scores on a knowledge assessment of evolution than students who had followed the standard curriculum. After completing our curriculum, students who were originally provided only the standard curriculum were able to further increase their assessment scores to meet the gains made by students who were taught evolution only with EvolvingSTEM. Our results demonstrate the power of microbial evolution experiments to effectively teach concepts in population genetics and evolution while also providing valuable experience in microbiology. Furthermore, EvolvingSTEM can serve as an instructional foundation of other life science topics. For example, further investigations by students could identify the genetic mutations (using inexpensive whole-genome sequencing, i.e. (Cooper 2018)) that underlie the adaptive mutant phenotypes, supporting a greater understanding of inheritance and trait variation (NGSS HS-LS3). Previous research in our lab indicates that many commonly identified mutations are found in the *wsp* (wrinkly spreader phenotype) gene cluster (Cooper et al. 2014; Gloag et al. 2018), which coordinates bacterial surface recognition with increased biofilm production (Hickman et al. 2005). Students are likely to identify *wsp* mutants in their classroom experiments and can therefore connect how changes in DNA can result in changes in protein structure and intracellular signaling that lead to increased biofilm production and changes to colony morphology, supporting a greater understanding of DNA, protein structure, and cellular function (NGSS HS-LS1). Furthermore, the bacterial adaptations are in response to environmental changes that provide new niches, supporting a greater understanding of interdependent relationships in ecosystems (NGSS HS-LS2). Classroom experiments that build upon the core evolution study can therefore span much of the NGSS-recommended introductory biology curriculum and have been adapted to cover more advanced topics for Advanced Placement (AP) Biology as well as to early biology courses in community colleges or four-year colleges.

This study was limited to one school and two teachers from a suburban public school in New Hampshire, which naturally raises the question of its efficacy in other settings. However, since the program launch and assessments reported here, EvolvingSTEM has expanded to be offered in 13 high schools in four different US states with continued growth. These schools range from independent private schools, to suburban public schools, to urban public and magnet high schools, and the classes include introductory “academic” and honors biology, upper-level biotechnology, and AP biology. The core experimental protocol described here has been shown to be robust to different class schedules and student populations, provided that the classroom has the laboratory resources detailed in Supplemental File 1, including the capacity to prepare sterile growth media either onsite or through a partner laboratory. Additional assessments of learning and motivation towards STEM subjects are ongoing in these schools, but informal teacher and student feedback has been overwhelmingly positive.

## Summary

EvolvingSTEM is an engaging, inquiry-based curriculum that provides students with a hands-on approach to visualize evolutionary change occurring in real time. It also can be delivered at a low cost per student (<$5 in consumables) and is therefore potentially suitable for broad distribution. Our curriculum provides students with the tools to understand evolutionary concepts and to apply their knowledge to other areas of life science and medicine. For example, students can make a direct link between the adaptive phenotypes they see in the classroom for increased biofilm production and the nearly identical phenotypes seen in clinically relevant biofilm-associated bacterial infections. In addition, students are provided an introduction to microbiological techniques that have important applications for biotechnology. A particularly powerful aspect of our curriculum is its positive effect on teacher and student engagement. Teachers and students embark on the research experiment together, which provides a collaborative classroom environment where both have the opportunity for greater understanding and discovery. EvolvingSTEM has exceptional ability to improve scientific literacy and the promise of promoting broad acceptance of evolution as a central, unifying theory for life science.

## Acknowledgements

We thank Stephen Hale from the University of New Hampshire for assistance in designing the assessment and Caroline Turner for critical feedback. This research was conducted according to IRB protocol #5937 from the University of New Hampshire supported by NSF CAREER DEB-0845851, NASA CAN-7NNA15BB04A, and discretionary funds from the University of Pittsburgh, School of Medicine to VSC.

## Supplemental Files

**Sup. 1: Materials Needed and Media Recipes**

**Sup. 2: EvolvingSTEM Experimental Protocol**

**Sup. 3: Curriculum Overview**

**Sup. 4: Pre and Post Lab Questions**

**Sup. 5: Student Test and Grading Rubric**

## Materials Needed per Classroom

### Materials for entire classroom

- Gloves (will need at least 6 pairs per student)
- Spray bottle of 70% Ethanol (to clean benchtop)
- Spray bottle of 10% Bleach (to decontaminate bacterial cultures and plates)
- Orbital shaker
- Incubator
- Serological pipettes and Pipette aid (to prefill tubes with media and PBS)
- *Pseudomonas fluorescens* SBW25 colonies streaked on ½ Tsoy-Agar plates (need to have 4 distinct colonies for each student group)
- Autoclave (to sterilize reusable materials and media)
- Dissecting microscope (not required, but helpful to visualize colonies)

### Materials for each student group

- Bunsen burner
- Vortex
- Micropipettes and tips: p200 and p1000
- Forceps: 1 pair
- Metal inoculation loop: 1
- Glass spreader beads
- Glass culture tubes (15mL): 36
- Small glass tubes (5mL): 8
- 5 and 15 mL tube racks
- White beads: 9
- Black beads: 3
- Queen’s B Media: 82mL
- PBS: 102mL
- Tsoy-agar plates: 12

### Media Recipes

#### 1L Queen’s B Media

- 20g Proteose Peptone No. 3
- 1.5g K_2_HPO_4_ (Potassium Phosphate Dibasic)
- 25mL Glycerol
- 970mL Water
  1. Autoclave for 45 minutes
  2. Allow to cool to room temperature
  3. Add 6mL of 1M MgSO_4_ (Magnesium Sulfate) stock

##### 250mL 1M MgSO_4_ Stock

- 30g MgSO_4_ (anhydrous) *or*
- 61.6g MgSO_4_ (heptahydrate)
- 250mL Water
  1. Combine salts and water
  2. Autoclave for 30-45 minutes

#### 1L PBS

- 7.65g NaCl
- 0.72g Na_2_HPO_4_ (Sodium Phosphate Dibasic, anhydrous)
- 0.21g KH_2_PO_4_ (Potassium Phosphate Monobasic)
- 1L Water
  1. Combine salts and water
  2. Autoclave for 45 minutes

#### 1L ½ Strength Tsoy-Agar (makes approximately 50 Plates)

- 15g Tsoy
- 15g Agar
- 1L Water
  1. Autoclave for 45 minutes
  2. Pour plates while still hot (so agar does not harden)
  3. Allow to solidify overnight before using

## EvolvingSTEM Pseudomonas fluorescens Experimental Evolution Protocol

*You are about to embark on a journey through a world that you might be unfamiliar with; one filled with odd instruments that you will use to study oddly shaped slimy bacterial colonies and neon yellow biofilm-coated test tubes. Over the course of the next few weeks you will be taking care of bacterial cultures, and your ordinary looking colonies will evolve to produce distinct mutants that have adapted to inhabit different parts of a test tube.*

### ALWAYS REMEMBER

Proper **aseptic technique** is a very important part of microbiology! All tubes, beads, and media have been sterilized in an **autoclave** prior to use in these experiments. When tubes were prepared, media was always distributed using sterile pipettes, and sterile beads were added using forceps that have been heated over a flame until *“red-hot”* to prevent contamination.

### USEFUL TERMS

**Aseptic Technique** – a sterile set of practices and procedures performed to minimize contamination by other bacteria.

**Autoclave** – a strong, heated container that reaches high temperature and pressure to sterilize equipment and media.

### SAFETY FIRST!

**You will be working with an open flame during this experiment.** Always be aware of your surroundings to ensure that you do not burn yourself or start a fire. Be sure to know the location of the closest emergency shower and fire extinguisher in case an accident does occur.

**Always treat the bacteria you will be working with as potential pathogens** (even though *P. fluorescens* is harmless to humans!). Be sure to disinfect your work stations and waste materials with 10% bleach and always follow general safe lab procedures, including tying back long hair, washing your hands at the end of the lab activity, and wearing gloves and lab coats.

The following links provide excellent information on: General Lab Safety: https://www.youtube.com/watch?v=MEIXRLcC6RA&vl=en Safely Working with Microorganisms: https://www.sciencebuddies.org/science-fair-projects/references/microorganisms-safety

### DAY 1 (MONDAY): PRECONDITIONING YOUR BACTERIAL CULTURE

*Before the bacteria got to your classroom, they had been stored in a freezer for a long period of time at −80*° *Celsius (−112*° *Fahrenheit). Before we can continue with our experiment, we want to ensure that the bacteria are used to being out of the freezer so they are performing at their prime. In order to do so, we give them time to get acclimated to their new environmental conditions – this day is known as our “Preconditioning Day”.*

#### NECESSARY MATERIALS

- Inoculation loops
- *Pseudomonas fluorescens* SBW25 colonies (on agar plate)
- 3 Large glass culture tubes containing: 5 mL Queen’s B Medium (QB) 1 **white** polystyrene bead
- 1 Large glass culture tube containing: 5 mL Queen’s B Medium (QB)

**Figure 1:**
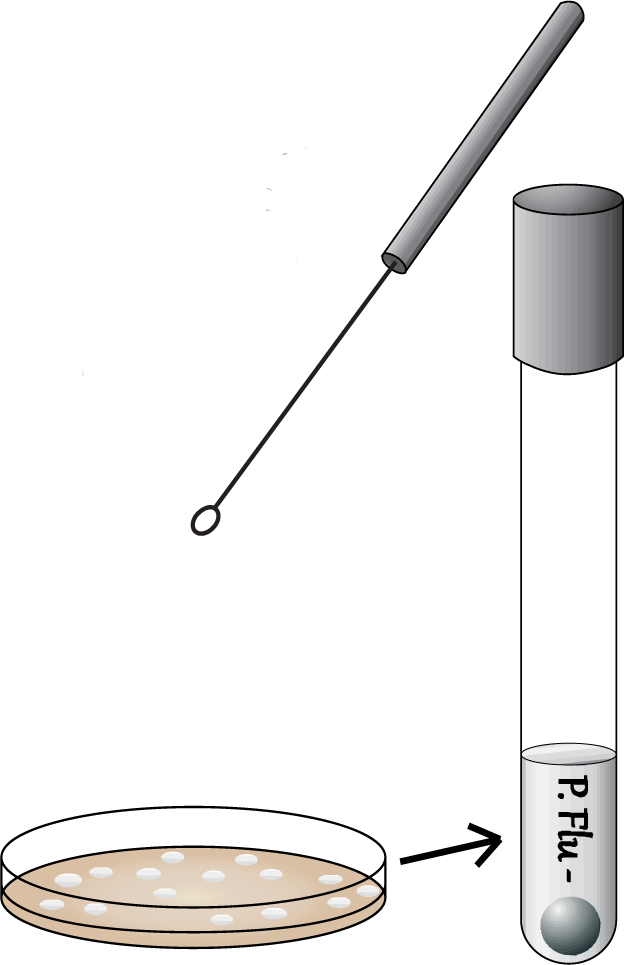
Use a sterile loop to inoculate each tube with a single colony.

#### PROCEDURE

1. Use an inoculation loop to transfer a **single** isolated *P. fluorescens* colony to a **single** culture tube.
2. Repeat until you have inoculated all four tubes: “1”, “2”, “3”, “C”. **BE SURE TO USE A NEW COLONY TO INOCULATE EACH TUBE!**
3. Incubate the culture tubes on a rotating shaker at 28°C until your next class.

### DAY 2 (TUESDAY): BEAD TRANSFER AND PLATING

#### NECESSARY MATERIALS

- Metal forceps
- 4 Small glass tubes containing 1 mL QB
- Vortex
- A p200 and p1000 pipette
- 3 Large glass evolution tubes containing: 4.5 mL QB 1 white polystyrene bead
- 1 Large glass control tube containing: 4.5 mL QB
- 8 Large glass culture tubes containing 5 mL Phosphate Buffered Saline (PBS)
- 4 ½ Strength Tsoy-Agar plates with small glass beads

#### PROCEDURE

1. Label the large glass culture tubes and agar plates in an identifiable manner.
2. Flame sterilize the forceps and allow them cool for 30 seconds. *After flaming the forceps they must not touch anything else, or they are no longer considered to be sterile!*
3. **For each evolution culture:** Pour the contents of the culture tube into its metal cap, and then use sterile forceps to transfer ONLY the bead to the corresponding small QB tube. *It is possible that you may hear a sizzle; this is normal and just means that the forceps are still hot from sterilization. Allow them to cool until you no longer hear a sizzle before you touch the polystyrene bead.*

**Figure 2:**
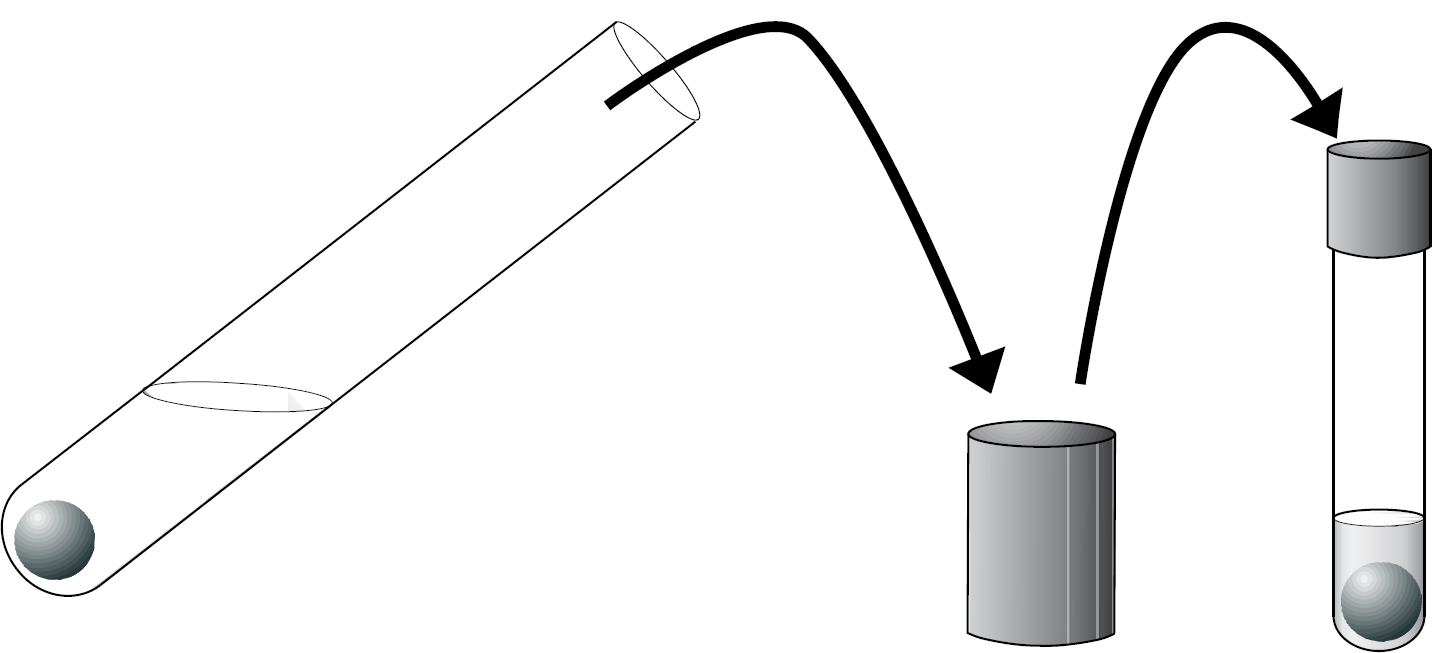
The preconditioning culture (left) is poured into its cap, and then sterile forceps are used to move only the bead to the small QB tube (right). Vortex the small QB tube for at least 45 seconds to remove biofilm from the bead. **For the control tube:** Briefly swirl the control culture tube, and then use a **p200** pipette to transfer 50 μl of the culture to the small QB tube. Briefly vortex the small QB tube.

#### Perform the following steps for all evolution and control cultures

4. Use a **p1000** pipette to transfer 500 μl from the small QB tube to the large QB tube. Briefly vortex to mix.
5. Use a **p200** pipette to transfer 50 μl from the large QB tube to a PBS tube (10^−2^ dilution). Briefly vortex to mix.
6. Use a **p200** pipette to transfer 50 μl from the 10^−2^ tube to a new PBS tube (10^−4^ dilution). Briefly vortex to mix.
7. Use a **p200** pipette to transfer 100 μl of the 10^−4^ dilution to an agar plate.
8. Shake the plate with the lid on top using the glass beads to spread the liquid culture. Remove the glass beads by turning the plate upside down and dumping the beads from the lid into the container provided.
9. Incubate the culture tubes on a rotating shaker at 28°C until your next class. Incubate the plates, **upside down**, at 28°C until your next class.

**Figure 3:**
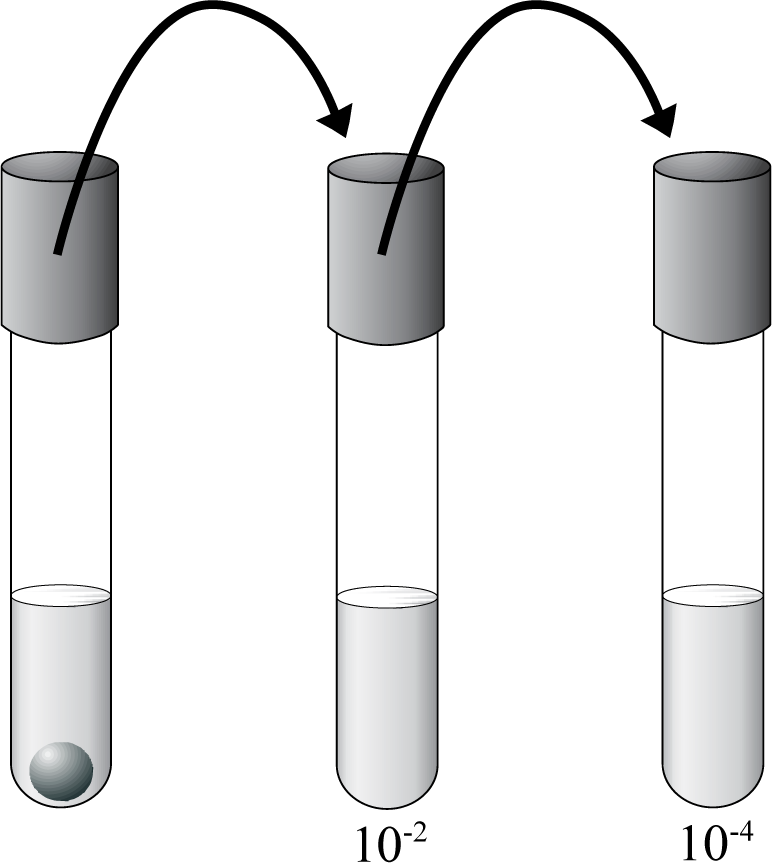
Serial dilution from the evolution tube (left) into PBS.

### DAY 3 (WEDNESDAY): BEAD TRANSFER

*The millions of cells that you added to your tube will quickly grow to become billions. It doesn’t take long before the bacteria consume the food and nutrients provided by the media inside of the test tube. In order to make sure that the bacteria continue to survive, we have to transfer a small number into a new tube. In the case of our experimental cultures, we transfer only the bacteria that are good at forming biofilm and have thus successfully stuck to the bead.*

#### NECESSARY MATERIALS

- Metal forceps
- Vortex
- p200 pipette
- 3 Large glass evolution tubes containing: 5ml QB 1 **black** polystyrene bead
- 1 Large glass control tube containing: 5ml QB

#### PROCEDURE

1. Flame sterilize and cool the forceps.
2. **For each evolution culture:** Pour the contents of the culture tube into its metal cap, and then use sterile forceps to transfer the **white bead** to a new evolution tube with fresh media and a **black bead.** **For the control culture:** Briefly swirl the control culture tube, and then use the **p200** pipette to transfer 50 *μ*l of the culture to the new control tube.
3. Incubate the culture tubes on a rotating shaker at 28°C until your next class.

### DAY 4 (THURSDAY): BEAD TRANSFER

*You may have noticed that your incubated test tubes now contain both a white and a black bead. Today, you are transferring your black bead to a new tube containing fresh media and a white bead. Over time, some of the bacteria from the black bead will detach and re-adhere to the surface of the white bead.*

#### NECESSARY MATERIALS

- Metal forceps
- Vortex
- p200 pipette
- 3 Large glass evolution tubes containing: 5ml QB 1 **white** polystyrene bead
- 1 Large glass control tube containing: 5ml QB

#### PROCEDURE

1. Flame sterilize and cool the forceps.
2. **For the evolution tubes:** Pour the contents of the culture tube into its metal cap, and then use sterile forceps to transfer the **black bead** to the new corresponding evolution tube with fresh media and a **white bead.** **For the control tube:** Briefly swirl the control tube culture, and then use the **p200** pipette to transfer 50 μl of the culture to the new control tube.
3. Incubate the culture tubes on a rotating shaker at 28°C until your next class.

### DAY 5 (FRIDAY): FINAL PLATING

#### NECESSARY MATERIALS

- Metal forceps
- Vortex
- A p200 and p1000 pipette
- 4 Small glass tubes containing 1 mL Phosphate Buffered Saline (PBS)
- 8 Large glass culture tubes containing 5 mL Phosphate Buffered Saline (PBS)
- 4 Large glass culture tubes containing 4.5 mL Phosphate Buffered Saline (PBS)
- 8 ½ Strength Tsoy-Agar plates with glass beads

#### PROCEDURE

1. Flame sterilize and cool the forceps.
2. **For each evolution tube:** Pour the contents of the culture tube into its metal cap, and then use sterile forceps to transfer the **black bead** to the small glass tube with PBS. Vortex the small PBS tubes for at least 45 seconds to remove cells from the bead. **For the control tube:** Briefly swirl the control tube culture, and then use a **p200** pipette to transfer 50 μl of the culture to a small glass tube with PBS. Briefly vortex the small PBS tube.

#### Perform the following steps for all evolution and control cultures

3. Use a **p200** pipette to transfer 50 μl from the small PBS tube to a 5mL PBS tube (10^−2^ dilution). Briefly vortex to mix.
4. Use a **p200** pipette to transfer 50 μl from the 10^−2^ tube to a new 5mL PBS tube (10^−4^ dilution). Briefly vortex to mix.
5. Use a **p1000** pipette to transfer 500 μl of the 10^−4^ tube to the 4.5mL PBS tube (10^−5^ dilution). Briefly vortex to mix.
6. Use a **p200** pipette to transfer 100 μl of the 10^−4^ and 10^−5^ dilution tubes to agar plates
7. Shake the plates with their lids on top using the glass beads to spread the liquid culture. Remove the glass beads by turning the plates upside down and dumping the beads from the lid into the container provided.
**8.** Incubate the plates, **upside down,** at 28°C until your next class.

### DAY 6 (MONDAY): COLONY EXAMINATION

#### NECESSARY MATERIALS

- Dissecting microscope

#### PROCEDURE

Closely examine colony morphology:

- Do all colonies look exactly the same as those plated last Monday?
- If not, how many are different?
- Describe the following for each colony type:

- Size – large, medium, or small
- Texture – smooth or rough
- Color
- Shape

Use the following chart to help describe changes in colony appearance:

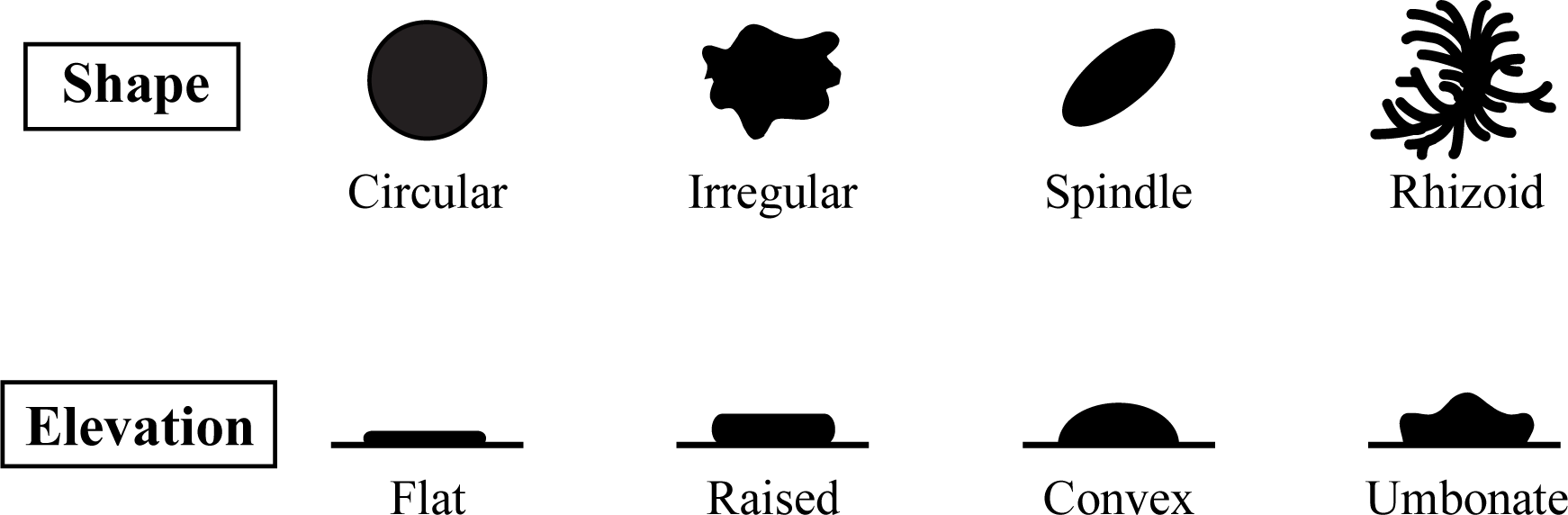

## EvolvingSTEM Curriculum Overview

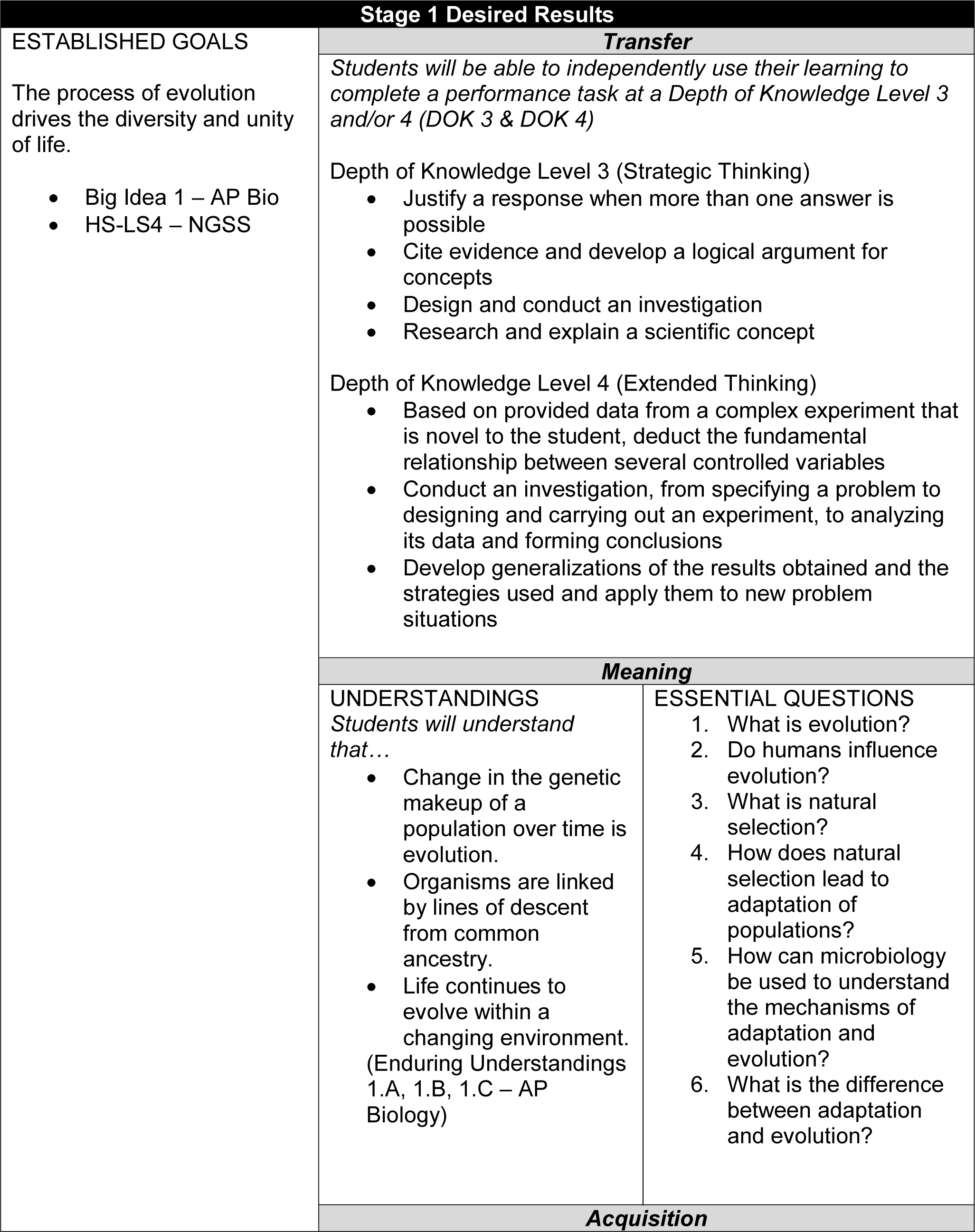
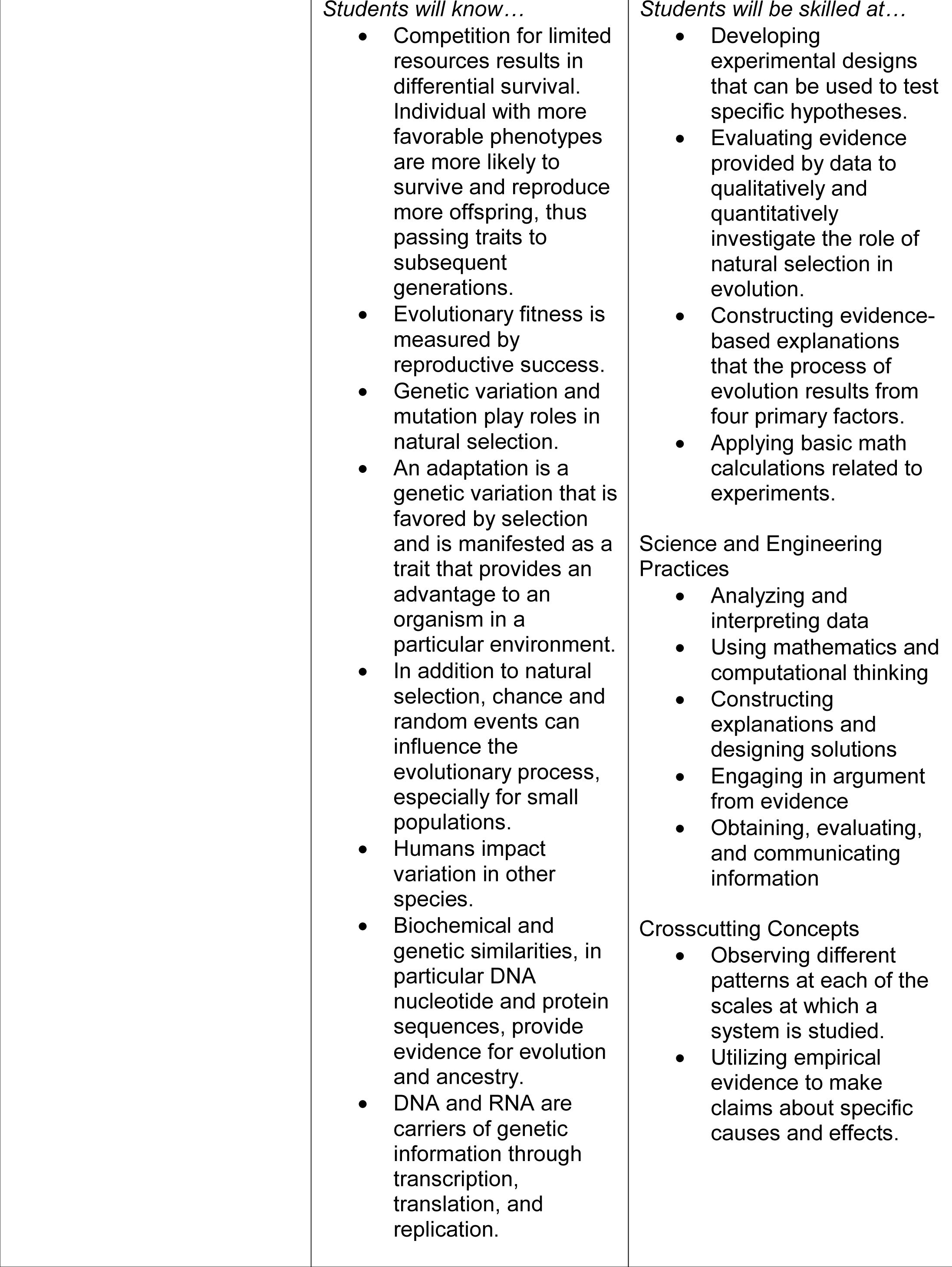
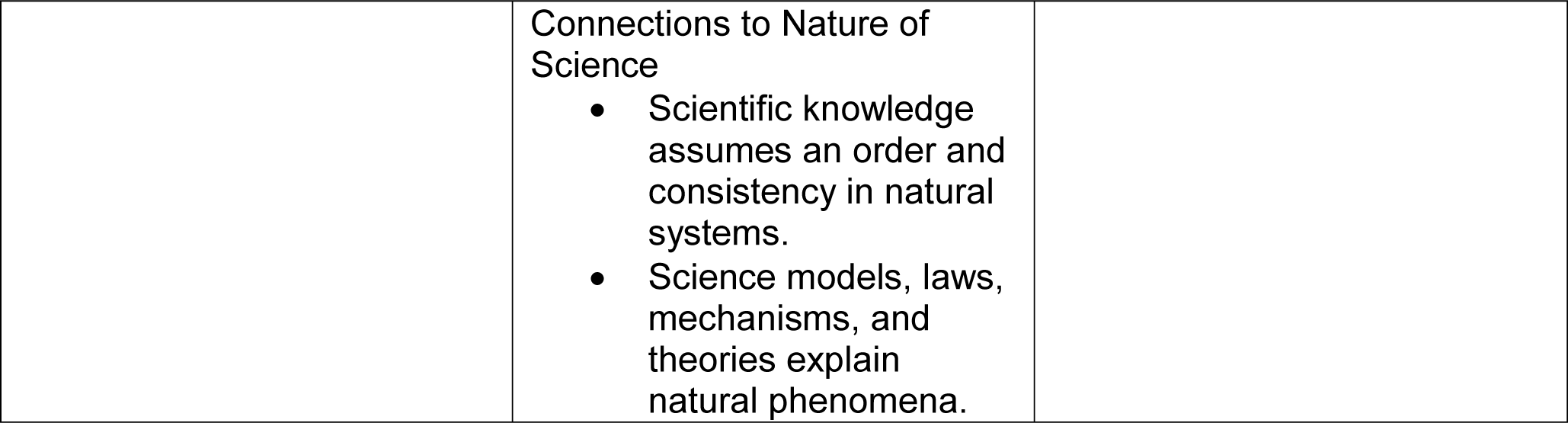

| Stage 2 – Evidence |  |
| --- | --- |
| Evaluative Criteria | Assessment Evidence |
| Project Rubric<br>Peer evaluation<br>Self-evaluation<br>Presentation with rubric<br>science showcase<br>school board meeting<br>conference<br>step up day<br>vertical team meeting<br>middle schools/other high schools<br>in class | **PERFORMANCE TASK(S):**<br>Given specific experimental conditions, predict the outcome of bacterial growth supported by existing research. (2-3 days).<br>Student-designed projects to change experimental conditions such as:<br>Intermediate levels of disturbance – shaking, non-shaking, mix between shaking and non-shaking.<br>Altering nutrient levels<br>Oxygen availability<br>Balance between oxygen and nutrients – short and fat or long and thin microcosms<br>Changing incubation conditions (2 weeks) |
| Journal grade with rubric<br>Daily participation grade (5 point/day)<br>Numerical test and quiz grades<br>Teacher and peer feedback<br>Self-reflection | **OTHER EVIDENCE:**<br>Lab Journal – daily entries<br>Class Starter and Exit Ticket questions – review previous day, lab protocol or technique, address misconceptions, standardized test release questions<br>Lab participation – daily grade<br>Probes as formative assessment to address misconceptions, identify areas of reteaching, and document student growth.<br>Quizzes and tests - document learning of content knowledge and lab skills |

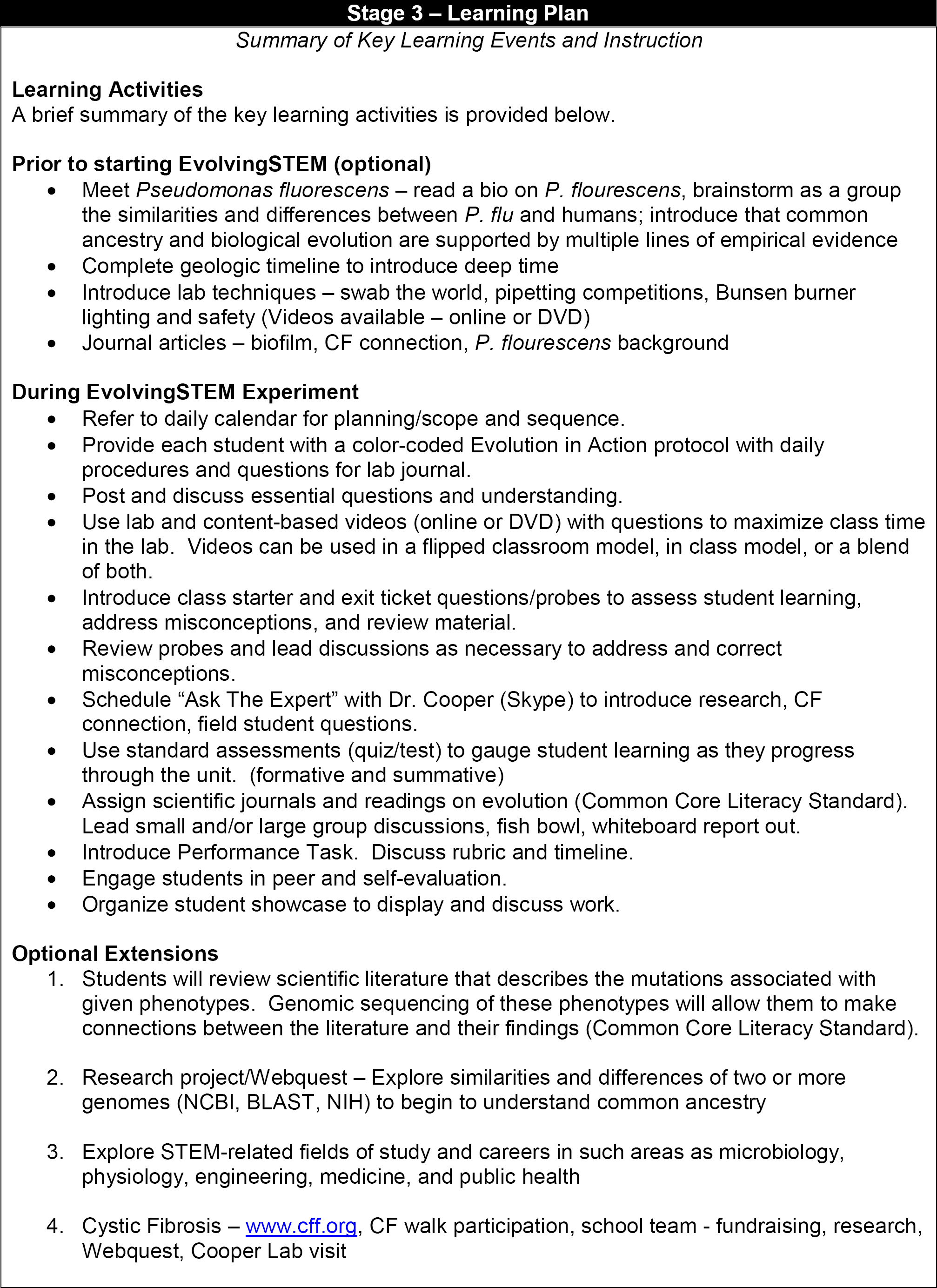
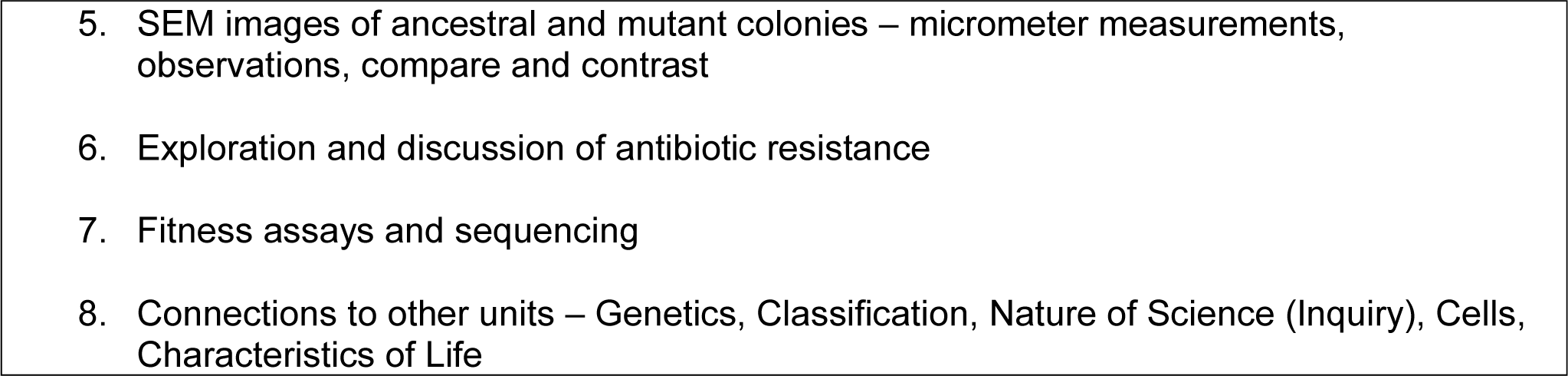

## PRE-LAB QUESTIONS: DAY 1

1. In one sentence briefly describe the purpose of the “pre-conditioning” step that will be carried out on Day 1 of your experiment? How many colonies are used to inoculate one test tube containing media and a white bead?
2. Suzie has just used a sterilized inoculating loop to obtain a single isolated colony of bacteria, which she then transferred into her test tube containing fresh media and a white bead. After she has put the cap of her newly inoculated test tube back on, she grabs the petri dish and goes to grab another colony. Before she can touch the inoculating loop to the petri dish, Larry stops her and tells her that she is doing it wrong. Which student is correct in this case, and why?

## POST-LAB QUESTIONS: DAY 1

1. What is another name of an error that is introduced during the process of replication that results in a new DNA sequence? What is the end product of this newly formed DNA sequence in comparison to the original DNA sequence?
2. What are the two possible forces that can act on mutations that occur in DNA sequences? Describe in detail the difference between these two different forces.
3. Describe the characteristics of bacteria that make them advantageous when studying evolution?
4. Summarize the different stages that occur throughout the biofilm lifecycle. How does this relate to the bead transfer model that is used in the experiments?

## PRE-LAB QUESTIONS: DAY 2

1. In one sentence briefly summarize the process of serial dilutions. What is happening to the overall population size of the bacteria as you carry out these dilutions and what is achieved by completing them?
2. Draw the series of steps that are required to complete a serial dilution on Day 2. Include the amount of liquid that is being transferred, the amount of liquid that is in the dilution tube, and the dilution that is achieved with each step. Circle the dilution(s) that will be plated on Day 2.
3. In one sentence briefly summarize the process of plating a bacterial culture. What is achieved by plating, and why is it incredibly important to ensure that you are plating on the agar side of the plate?
4. Once the bead has been transferred from the large glass evolution tube to the small glass tube containing 1 mL of Queen’s B media, how long should the small glass tube be vortexed for? What is the purpose of vortexing?
5. In general, the large media tubes will contain 5 mL of media; however, on Day 2 the large glass evolution tube only contains 4.5 mL of media. Can you explain why this is the case?

## POST-LAB QUESTIONS: DAY 2

1. Why is it important to transfer the bacteria every 24 hours? Draw a graph that illustrates the growth of a bacterial culture. Make sure to label your axes!
2. Provide a detailed hypothesis that describes what you think might occur in your test tube over the next 24 hours when your bacteria from inside the test tube are adhering to the new bead. Try to use the following vocabulary in your predictions: planktonic, biofilm, and overproduction and polystyrene bead.

## PRE-LAB QUESTIONS: DAY 3

1. What is the color of the old bead that is being transferred from the 24-hour large glass evolution tube? What is the color of the new bead that is in the new large glass evolution tube?
2. Why is it important to disrupt the bead as little as possible during your daily bead transfer?
3. Describe what your test tube looked like on Day 2 when you put it in the incubator following your transfer. What do you think it will look like on Day 3 when you remove it from the incubator? Describe the amount of biofilm that is seen on the sides of the tube, the type of biofilm that is seen, and the color of the liquid media.

## POST-LAB QUESTIONS: DAY 3

1. Describe in detail the three different types of mutations that can occur and the possible effect of that given type of mutation. Which two types of mutations are generally more common, and which is the least likely to occur?
2. What is being accomplished by transferring only the bacteria that have successfully attached to the bead? What is this type of selection called?
3. Provide a detailed hypothesis that describes what you believe might occur in your test tube over the next 24 hours when your bacteria are detaching from the old bead and adhering to the new bead. Use the following vocabulary in your predictions: overproduction, planktonic, biofilm, exponential, polystyrene bead, and mutation (both beneficial and neutral).

## PRE-LAB QUESTIONS: DAY 4

1. What is the color of the old bead that is being transferred from the 24-hour large glass evolution tube?
2. Describe what your test tube looked like when you put it in the incubator following your transfer on Day 3. What do you think it will look like on Day 4 when you remove it from the incubator? Describe the amount of biofilm that is seen on the sides of the tube, the type of biofilm that is seen, and the color of the liquid media.

## POST-LAB QUESTIONS: DAY 4

1. It is possible that when you removed your tubes today, only one of them has significantly more biofilm on the sides of the tubes and has a neon culture. As we discussed, this is a possible indication that you have a beneficial mutation in your population. Can you provide an explanation for why only one of your four replicates looks like this if you started with identical bacteria at the beginning of your experiment?
2. If we were to impose a greater force of artificial selection on the bacteria that we are studying, would it increase the number of mutations that we see in our experiment? Why or why not?
3. We already know that bacteria grow at an incredibly fast rate and can potentially overproduce, causing them to produce more bacteria inside the test tube than can survive. This over production leads to another phenomenon, which is another point of Darwin’s Theory of Evolution by Natural Selection. Explain how this is occurring inside of the test tube and how it relates to overproduction.
4. Provide a detailed hypothesis that describes what you believe might occur in your test tube over the next 24 hours when your bacteria are detaching from the old bead and adhering to the new bead. Use the following vocabulary in your predictions: overproduction, planktonic, biofilm, exponential, competition, mutation (both beneficial and neutral), resources, space, polystyrene bead, nutrients, frequency, niche, and heritable genetic variation.

## PRE-LAB QUESTIONS: DAY 5

1. What is the color of the bead that is going to be plated? Why isn’t it necessary to transfer the other bead to a new evolution tube containing fresh media and an oppositely marked bead?
2. Draw the series of steps that are required to complete a serial dilution on Day 5. Include the amount of liquid that is being transferred, the amount of liquid that is in the dilution tube, and the dilution that is achieved with each step. Circle the dilution(s) that will be plated on Day 5.
3. Provide a detailed hypothesis as to why you believe it is necessary to dilute one step further on Day 5 than on Day 2.
4. Describe what your test tube looked like when you put it in the incubator following your transfer on Day 4. What do you think it will look like on Day 5 when you remove it from the incubator? Describe the amount of biofilm that is seen on the sides of the tube, the type of biofilm that is seen, and the color of the liquid media.

## POST-LAB QUESTIONS: DAY 5

1. Did one of your tubes change drastically from Day 4? If you observed a change in one of your tubes, explain the differences that you are seeing. In addition, provide possible explanations as to why you have yet to see a change in your other replicates.
2. Discuss how mutations increase in frequency over time. You may draw a graph below to illustrate this process.
3. When you removed your tube from the incubator on Day 2, it had not appeared to change greatly from when you started your experiment. When you removed your tube from the incubator on Day 5, the culture was neon yellow. You are sure that when you plate today, you will definitely have mutants on your plate. Your group member also states that it is possible that you have mutants on your Day 2 plates. Is he/she correct? Discuss the potential results that may be observed.

## PRE-LAB QUESTIONS: DAY 6

1. Predict what your Day 5 colonies will look like when you view them in the lab. How will they look different from the colonies you plated on Day 2?
2. As we discussed previously, it is possible that you may see multiple phenotypes on your agar plate during the course of your evolutions. Provide a hypothesis that might explain the role that each of these mutants is playing in the community.

## POST-LAB QUESTIONS: DAY 6

1. Explain how two mutants with distinct phenotypes can inhabit the same test tube simultaneously. Be sure to incorporate the importance of an ecological niche in your answer.

1. Now that you have completed your evolution experiment, do you believe that evolution is fast or slow? Provide an explanation to support your answer.
2. You now have all four pieces that are required to support Darwin’s Theory of Evolution by Natural Selection. Use all four to comprise an explanation that can support our example of microevolution that occurred in our test tube over the past week. Do you think that these same four points can be applied to a macro-evolutionary example?
3. Explain the difference between evolution and adaptation.

## STUDENT TEST

Student ID Number:___________________

Block Number:________________________

Teacher:_____________________________

Date:_________________________________

(Understanding)

From the groups of characteristics below, identify the best answer for describing evolution. (2 pts. each)

1. Rate of evolution:

a. Evolution does not happen, the rate of change in species is zero
b. Fast – taking place in just a few generations
c. Slow – taking thousands of generations or many thousands of years
d. Evolution can be either Fast or Slow
2. The fundamental source of genetic variation among individual organisms is:

a. Levels of nutrition that individuals receive
b. Random mutations in DNA sequences or chromosomes
c. Physical changes accumulated during an organism’s lifetime
d. Unexpected changes occurring during embryonic development
3. Amount of change in evolution:

a. Evolution occurs rapidly, with quick appearance of new traits
b. Evolution occurs at rates ranging from gradual to rapid
c. Evolution occurs gradually by the accumulation of small changes over time
d. Evolution does not happen so the amount of change is zero
4. Types of organisms that evolve:

a. Evolution does not happen in any type of organism
b. Evolution occurs in tiny organisms like bacteria and other single-celled species
c. Evolution occurs in large organisms like palm trees, crabs, snakes, and giraffes
d. Evolution occurs in all groups of organisms
5. Which example statement best describes evolutionary change?
  a. There are no observable changes in organisms over time
  b. A cat fed on a good diet grows larger than a cat fed on a poor diet
  c. A fair-skinned person tans during a summer
  d. Plants growing on a wet, lush island grow higher than plants on a dry, desert island
  e. The bill shape of birds changes because the hardness of the seeds they eat changes
6. For evolution to occur, which genetic characteristic must be present?
  a. There must be an even number of chromosomes
  b. Individual organisms in a population must appear different
  c. The differences in organisms must be capable to be passed to offspring
  d. All of the characteristics listed above must be present (Apply)
7. The New Mexico Whiptail (*Cnemidophorous neomexicanus)* is a parthenogenic lizard species from New Mexico and Arizona. The population consists entirely of females capable of laying viable eggs without fertilization. Many years ago, after receiving the necessary collecting permits, your teacher collected a single (1) New Mexico Whiptail and brought it to your school laboratory in NH. Your school laboratory is well-resourced with aquaria and mealworms, and everything needed to rear healthy whiptails. The first clutch yielded 10 offspring from the original whiptail and they were divided into two equal groups (A and B). Individuals in group A were marked by clipping the tip of the last digit on the left hind toe, and group B by the same procedure on the right hind toe. Note that the toe clipping has no effect on the ability of the lizard to survive in the environment, and was only used as a way to distinguish the two groups. Your teacher has been observing the mothers of the egg clutches, and as young whiptails hatch you clip the appropriate toe to assign them to the proper group of their mother. After 500 generations of living together with hatches and deaths occurring, the population size has grown to 340 whiptails with 272 in Group A, and 68 in Group B.

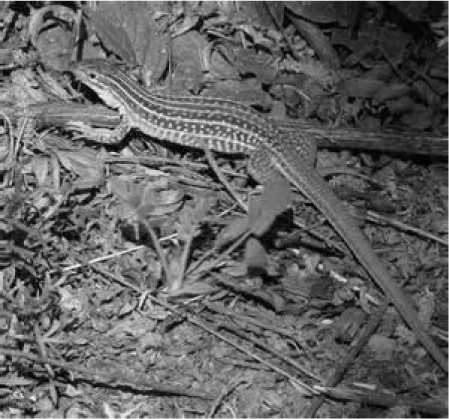 Count of Group A and B individual New Mexico Whiptails (*Cnemidophorous neomexicanus*) in the first generation (1) and in generation (500). Count does not include dead individuals, which were removed.

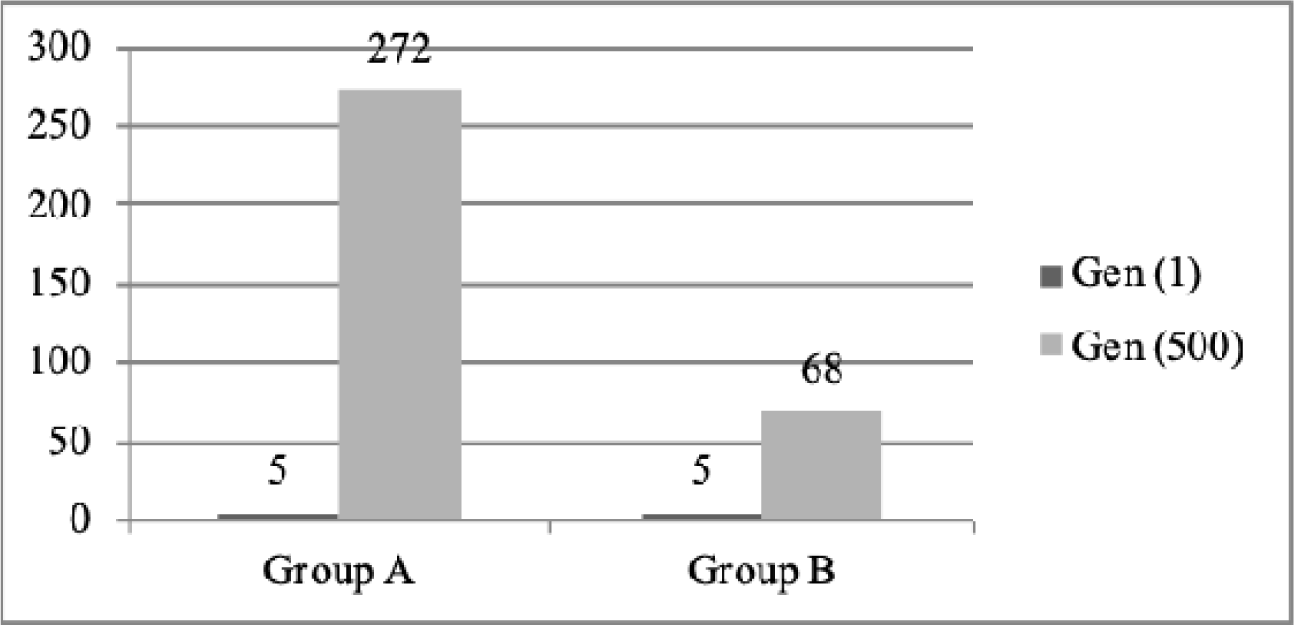

a. What is the percentage of Group A and Group B in the first generation of Whiptails? (2 points)
b. What is the percentage of Group A and Group B in the total population after 500 generations of hatches and deaths? (2 points)
c. Provide a possible evolutionary explanation for the shift in numbers of Group A and Group B individuals. (6 points) ________________________________________________________________________________ ________________________________________________________________________________ ________________________________________________________________________________ ________________________________________________________________________________ ________________________________________________________________________________ ________________________________________________________________________________ ________________________________________________________________________________ ________________________________________________________________________________ ________________________________________________________________________________ ________________________________________________________________________________ ________________________________________________________________________________ _____________
8. List and briefly describe the 4 key elements that produce evolutionary change (12 points, 1.5 points for each correct concept, and 1.5 points for each correct description). 1. 2. 3. 4. (Analyze)
9. You and your lab partner are given a test tube with a single living type of bacterium that grows in the water at room temperature. When you grow the organism in a petri dish, the bacterium only grows in circular-shaped colonies with clean, smooth edges. Then you grow your bacterium in the water but in a refrigerator. When you grow the organism from the refrigerator in a petri dish, you find circular-shaped colonies with clean, smooth edges, but also colonies with irregular-shapes and rough, jagged edges. You and your lab partner repeat this procedure, each time beginning with only the original bacterium. Each time you get the same result. Your lab partner exclaims, “The bacterium evolved!!” Is your lab partner correct? Why or why not? (16 points)

## Pre- and Post-Test Grading Rubric

|  | EXCEEDS<br>4 | MEETS<br>3 | APPROACHES<br>2 | BEGINS<br>1 | STRUGGLES<br>0 |
| --- | --- | --- | --- | --- | --- |
| Content Knowledge | Information on the topic is accurate. It addresses and extends beyond the questions raised in the prompt. Student incorporates outside examples that strongly help them defend their claims. Important vocabulary is used properly in the context of the response. It synthesizes competing ideas and says something new. | Information on the topic is accurate. It addresses the questions raised in the prompt. Important vocabulary is used, but not necessarily in the proper context. | Information on the topic is accurate, but does not address all questions raised in the prompt. Important vocabulary is not used. | Information on the topic is inaccurate or sparse. The response reports facts. | No response is supplied or the information on the topic is incorrect. |

